# ERStruct: A Python Package for Inferring the Number of Top Principal Components from Whole Genome Sequencing Data

**DOI:** 10.1101/2022.08.15.503962

**Authors:** Jinghan Yang, Yuyang Xu, Minhao Yao, Gao Wang, Zhonghua Liu

**Affiliations:** The University of Hong Kong; Columbia University

## Abstract

Large-scale multi-ethnic DNA sequencing data is increasingly available owing to decreasing cost of modern sequencing technologies. Inference of the population structure with such sequencing data is fundamentally important. However, the ultra-dimensionality and complicated linkage disequilibrium patterns across the whole genome make it challenging to infer population structure using traditional principal component analysis (PCA) based methods and software. We present the ERStruct Python Package, which enables the inference of population structure using whole-genome sequencing data. By leveraging parallel computing and GPU acceleration, our package achieves significant improvements in the speed of matrix operations for large-scale data. Additionally, our package features adaptive data splitting capabilities to facilitate computation on GPUs with limited memory. Our Python package ERStruct is an efficient and user-friendly tool for estimating the number of top informative PCs that capture population structure from whole genome sequencing data.

## 1 Introduction

With the fast development and decreasing cost of next generation sequencing technology, whole genome sequencing (WGS) data are increasingly available and hold the promise of discovering the genetic architecture of human traits and diseases. One fundamental question is to infer population structure from the WGS data, which is critically important in population genetics and genetic association studies (1; 2; 3).

PCA-based methods are prevalent in capturing the population structure from array-based genotype data (4; 5; 6). However, it has been challenging to determine the number of top PCs that can sufficiently capture the population structure in practice. A popular traditional method (5) does not perform well on sequencing data for two reasons: ultra-dimensionality (7) and linkage disequilibrium (5). To resolve those two practical issues on sequencing data, a novel method ERStruct based on eigenvalue ratios has been proposed (8) for whole genome sequencing data and a MATLAB toolbox has been developed. They showed a substantial improvement in accuracy and robustness when applying the ERStruct toolbox both on the HapMap 3 project (9) array-based data and the 1000 Genomes Project (10) sequencing data. Despite the great potential, we found that two issues may restrict the use of the ERStruct algorithm:

1. Although the ERStruct MATLAB toolbox provides a parallelization computing feature, its scalability is heavily restricted by the size of memory available in the working environment, slowing down the speed of data analysis.
2. The ERStruct algorithm was implemented as a MATLAB toolbox which is not freely available. In contrast, Python is open-sourced and free to use.

To improve the efficiency and accessibility of the ERStruct algorithm, we develop a new Python package implementing the same ERStruct algorithm. Our ERStruct Python implementation uses parallelization computing to accelerate simulations of GOE (Gaussian Orthogonal Ensemble matrix) matrices used in the ERStruct algorithm. In addition, the package provides optional GPU acceleration to boost the speed of large-scale data matrix operations while maintaining feasible memory usage for GPUs with limited Video Random Access Memory (VRAM). We applied the ERStruct Python package to the 1000 Genomes Project data to demonstrate the computationally efficient performance in the results section. Compared with the original MATLAB version, we achieved a similar time spent using our Python implementation using only CPU (Central Processing Unit) acceleration and significantly reduced time consumption and memory usage with GPU acceleration.

## 2 Algorithm

For self-sufficiency of the article, in this section we briefly describe the ERStruct algorithm from (8). Given a *n*-by-*p* genotype data matrix *C* that consists of *p* genetic markers from *n* individuals. Each of the entry *C*(*i, j*) takes the value from {0, 1, 2}, which represents the raw count of the minor alleles for the genetic marker *j* on the individual *i*. The ERStruct algorithm estimates the number of top informative PCs that capture the latent population structure. Suppose there are *K* different (latent) subpopulations in total, and the *i*th individual is the *l*th person in the *k*th subpopulation, the ERStruct model for the genetic markers of this *i*th individual is given by

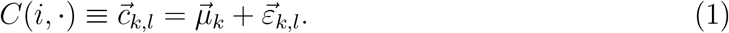

In this model, the *p*-dimensional vectors 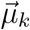 denote the *k*th subpopulation mean counts of minor alleles, and the vectors 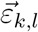 denote the individual noise vectors, which are independent and identically distributed with mean zeros and an arbitrary covariance matrix Σ.

We now summarize the key steps of the ERStruct. First, the data matrix *C* is normalized column by column into the new matrix *M* by

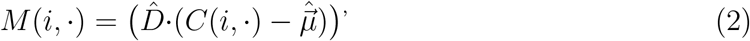

where

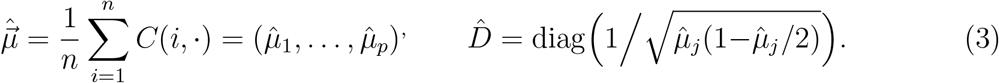

Next, we calculate the matrix *S*_*p*_ = *MM*^*/*^*p*, and compute its non-zero ordered sample eigenvalues ℓ_1_ = … = ℓ_*n*−1_ *>* 0, and obtain the corresponding sample eigenvalue ratios *r*_*i*_ = ℓ_*i*+1_*/*ℓ_*i*_, *i* = 1, …, *n* − 2.

According to the finite-rank perturbation theory (11; 12), under ERStruct model 1 and the following two assumptions:

1. ultra-high-dimensional asymptotic regime: *n* → ∞ and *n/p* → 0;
2. 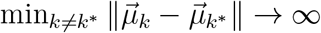

the sample eigenvalue ratios *r*_1_, …, *r*_*n*−2_ can be separated into two sets: the bulk and the spike. The bulk set contains the major part of eigenvalue ratios *r*_*K*_, …, *r*_*n*−2_, which will form a compact set and goes to 1 asymptotically. The spike set contains the remaining top *K* − 1 sample eigenvalue ratios *r*_1_, …, *r*_*K*−1_, which will converge (as *n* → ∞) to certain limits that are less than 1 and well-separated from the bulk set (see Fig. 1 for illustration).

**Figure 1:**
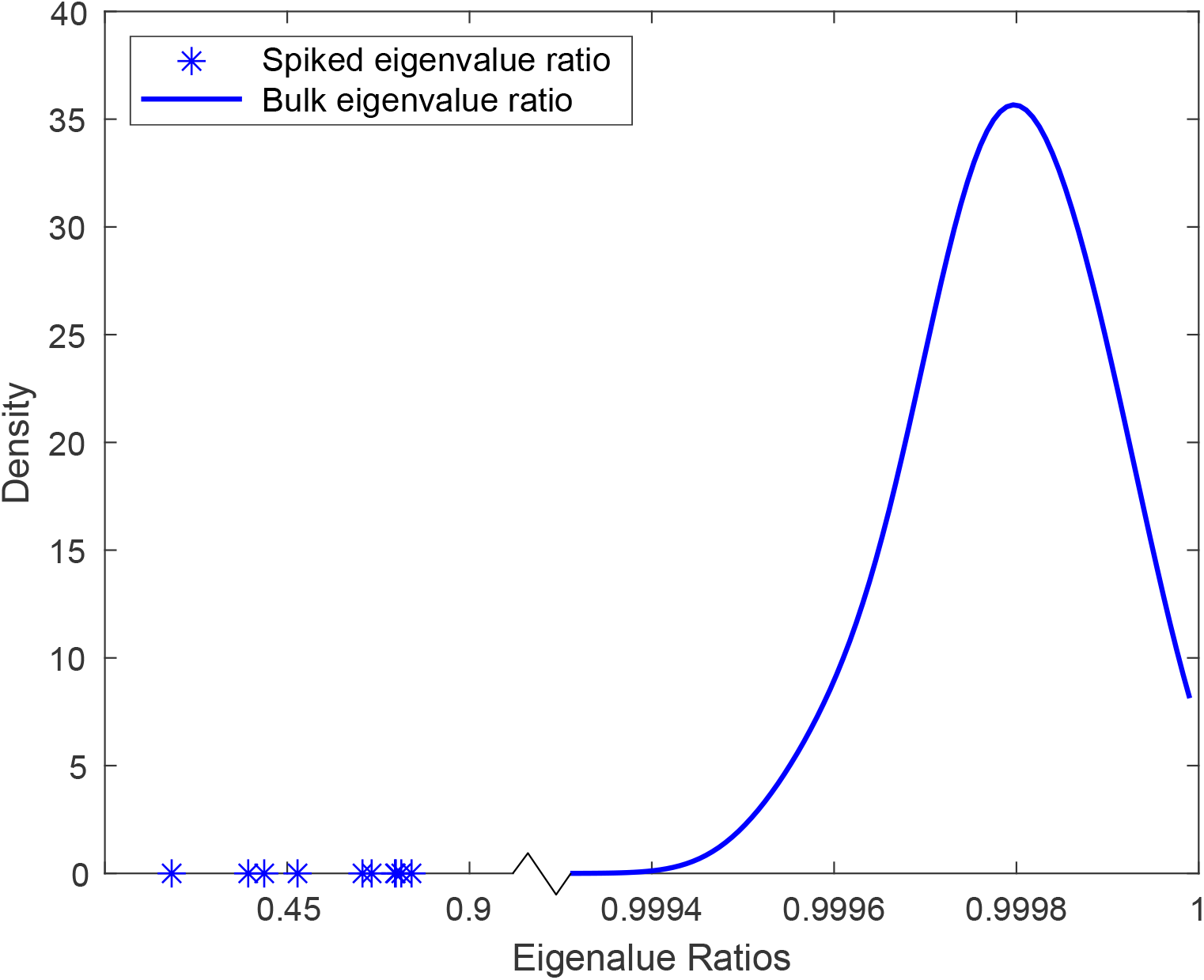
Illustration of a typical distribution of the eigenvalue ratios under the ERStruct model 1 (*K* = 12, *n* = 2500, *p* = 8000000).

The ERStruct algorithm estimates the number of top informative PCs *K* by estimating the number of spikes as follows,

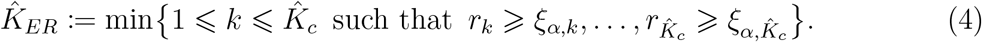

That is, the first index *k* such that the *k*th and the 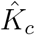 subsequent eigenvalue ratio 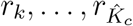 are respectively greater than their critical values 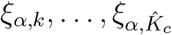, which are the lower *α* quantiles of the distribution of 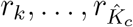. Note that there are two user-specified parameters in Eq. 4: the significance level *α* and the coarse estimator 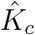. Based on the real data analysis results in (8), we recommend to use *α* = 0.001, and set the coarse estimate 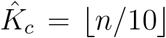 by default. Note that the coarse estimate should be generally larger than the true number of top PCs *K*, so as to avoid under-estimation of the original ER-based estimator proposed in (13),

To determine the critical value *ξ*_*α,k*_ for *r*_*k*_, the null distribution of *r*_*k*_ is needed and can be approximated by the distribution of the bulk eigenvalue ratio *r*_*K*_ when the sample size is *n* − *k* + 1 according to (12) (denotes as 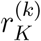). Applying the random matrix theory introduced in (14; 15; 16), the distribution of 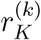 can be further approximated by

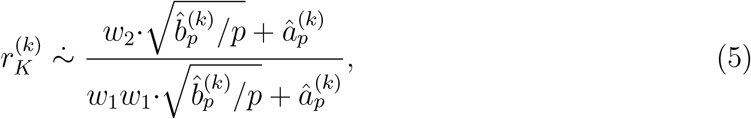

where the symbol 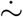 denotes the left and right sides asymptotically follow the same distribution, and

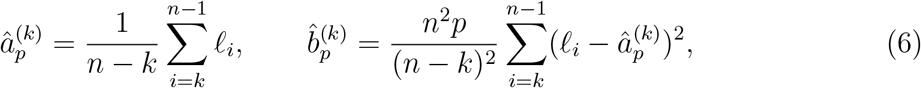

*w*_1_, *w*_2_ are the top two eigenvalues of a *n*-by-*n* GOE matrix (i.e., a square matrix with independent entries, where each diagonal entry follows *N* (0, 2) and each off-diagonal entry follows *N* (0, 1)).

Eqs. 5 and 6 link the distribution of (*w*_1_, *w*_2_) together to our target distribution of 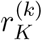, and thus the distribution of *r*_*k*_. In the ERStruct algorithm, Monte Carlo simulation is used to find out the empirical distribution of (*w*_1_, *w*_2_). Denote all the simulated replications as 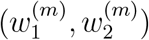, *m* = 1, …, *rep*, where *rep* is the number of replications. Then the empirical approximated distribution of *r*_*k*_ can be calculated by substituting (*w*_1_, *w*_2_) as 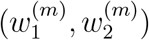 in Eq. 5, and sorting the results in ascending order (denotes as 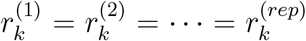). Finally, the critical value can be calculated by

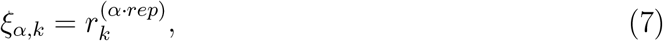

and the ERStruct estimator 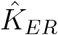 can be obtained by Eq. 4.

Note that to ensure validity, the replications *rep* should be in general greater than 1*/α*, the reciprocal of the input significance level. We recommend users to choose between 2*/α* to 5*/α* for the argument *rep*.

## 3 Implementation Details

For a given genotype data matrix as input, the ERStruct Python package estimates the number of top informative PCs that capture the latent population structure in three parts: GOE matrices simulation, calculating the eigenvalue ratios of the given data matrix, and estimating the number of top PCs. These parts correspond to `GOE.py`, `Eigens.py` and `TopPCs.py` files shown in Fig. 2, respectively. To boost the overall performance of the ERStruct algorithm, instead of the frequently used NumPy array processing framework, we choose to build up our ERStruct algorithm using PyTorch tensor and PyTorch functions. According to our experiments, PyTorch functions (e.g., `torch.nanmean`, `torch.nansum`, and `torch.linalg.eigvalsh`) are much faster than their NumPy equivalences (i.e., `numpy.nanmean`, `numpy.nansum`, and `numpy.linalg.eigvalsh`) for data processing in the ERStruct algorithm.

**Figure 2:**
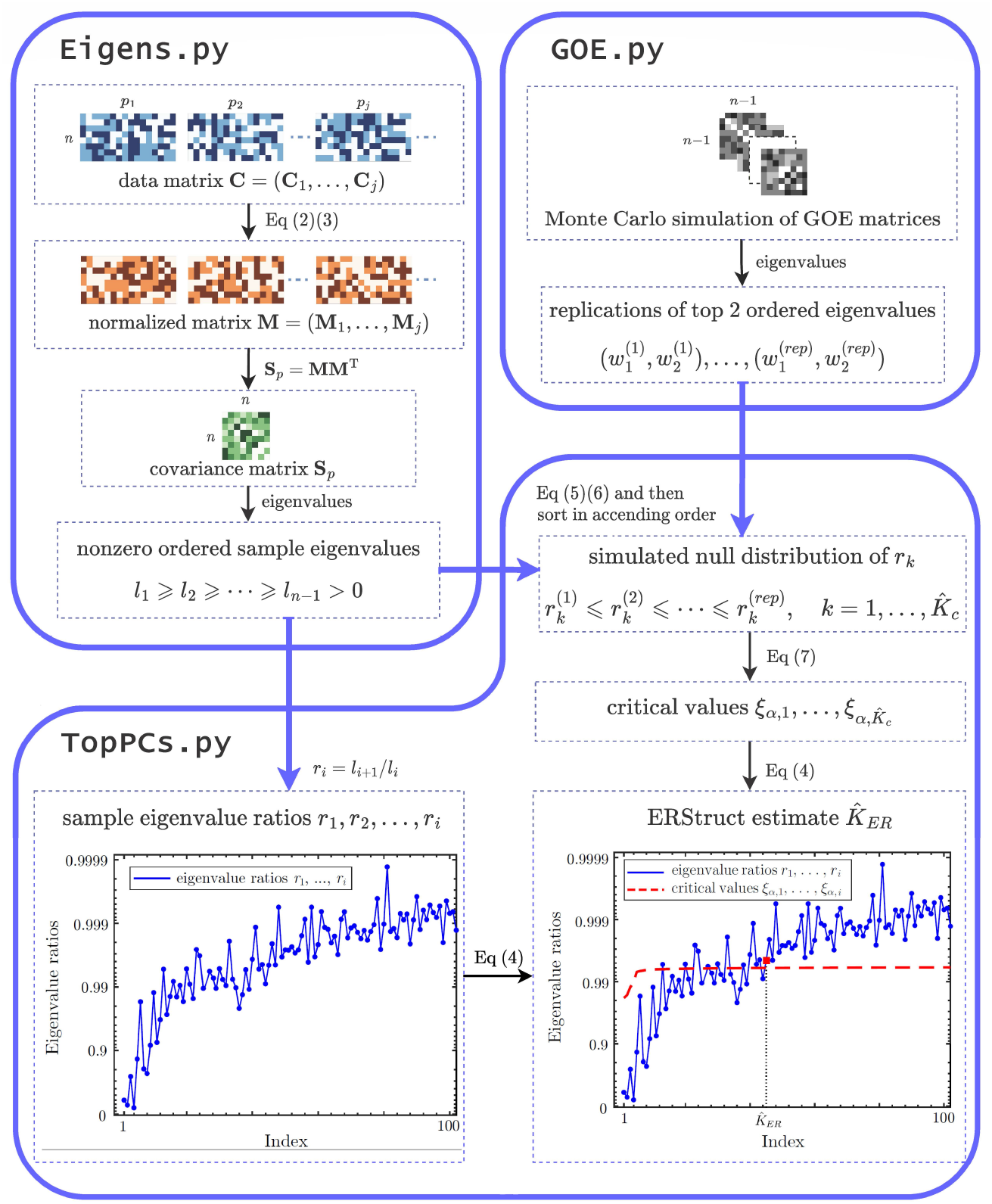
Flowchart that demonstrates the three parts of the ERStruct Python package for the whole genome sequencing data analysis. As an example we use the 1000 Genomes Project sequencing data set (10) in which genetic markers with MAF less than 5% are removed. The input real data are processed as in `Eigens.py` and then transmit to `TopPCs.py` to obtain the sample eigenvalue ratios *r*_*i*_ (as plotted on the lower left panel). While in `GOE.py`, GOE matrix simulation is carried out and then transmit to `TopPCs.py` to calculate the critical values 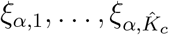. Finally, these critical values are used to infer the number of top principal components following Eq. 4 (as plotted in on the lower right panel).

### 3.1 Scalable Simulation of GOE Matrices

In `GOE.py`, Monte Carlo method is used in the ERStruct algorithm to obtain the null distribution of our proposed ERStruct test statistic, which starts by generating multiple replications of high-dimensional GOE matrices. A significant amount of computing resources is needed in this step, especially when the sample size of the experiment data is large. To efficiently simulate the high-dimensional GOE matrices, we have empirically tested different packages for parallelization computing on different scale GOE matrices, including Joblib, Multiprocessing, and Ray. Joblib is shown to have the most efficient and stable performance on our algorithm to apply parallelization computing for multicore. Without parallelization, the GOE matrix simulation takes 80.68 minutes on the function `GOE L12 sim` with sample size *n* = 2504 and the number of Monte Carlo replications *rep* = 5000, while using parallelization by Joblib on 15 cores CPUs, it takes only 6.46 minutes to finish the same job. Compared with the non-parallelization MATLAB version, we successfully decreased the computing time by 12.5 times.

### 3.2 Calculate the eigenvalue ratios of given matrix

In `Eigens.py`, we start from loading genotype data matrices *C*_*i*_ from NPY files as the input, where each row of matrix *C*_*i*_ represents the genetic markers of an individual. Multiple files input are allowed in this step because all of these large-scale data files are often too large to fit in the memory at the same time. In this case, users need to ensure that each data file alone can fit in the memory, and all of these data contain the same individuals in the same order. Also note that by default, our package will impute all the missing data by 0. To achieve a better performance potentially, users may perform other types of imputations beforehand.

The data matrix is then normalized according to Eqs. 2 and 3. In the next a few steps, sample covariance matrix *S*_*p*_ = *MM*^*/*^*p*, non-zero ordered eigenvalues ℓ_1_, …, ℓ_*n*−1_ and eigenvalue ratio *r*_*i*_ = ℓ_*i*+1_*/*ℓ_*i*_ (as shown in the lower left panel in Fig. 2) are calculated accordingly. Although the above matrix operations include only basic calculations, the data of interest are often massive, occupying a space of hundreds of gigabytes on disk, which may be expensive to process. In our ERStruct Python implementation, we accelerate the above computing steps by converting the CPU Tensor to a CUDA Tensor whenever GPU is available, taking the advantage of fast matrix operations in GPU. Limited VRAM capacity on many GPUs makes it difficult to process massive data matrices. However, our approach demonstrates the flexibility of data splitting as a solution to this challenge when working with large datasets on resource-constrained hardware. Our package automatically splits the input data into multiple sub-arrays using a CPU before transmitting it to the GPU. The size of the sub-arrays is determined based on the available VRAM capacity in the working environment. By adopting this approach, users can accelerate computations on large-scale sequencing data, even with a small VRAM GPU, while simultaneously reducing overall memory usage. Our approach highlights the potential of data splitting as an effective method for handling large datasets on hardware with limited resources. Users may use other popular genotype data formats like VCF (variant call format) and bgen files. A tutorial to convert VCF or bgen files to NPY files can be found in the ERStruct code repository.

### 3.3 Estimation of the number of top informative PCs

Finally, the sample eigenvalues ℓ_1_, …, ℓ_*n*−1_ from `Eigens.py` and the top two eigenvalues of simulated GOE matrices 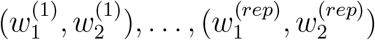 from `GOE.py` are transmitted together into `TopPCs.py`. Following Eqs. 4 to 7 in the ERStruct algorithm, the program output the estimation of the number of top informative PCs 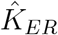 in the end (as shown in the lower right panel in Fig. 2).

Note that if the target estimator 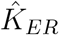 can not be found through Eq. 4, an error message will appear. This may happen if the data exhibit serious multicollinearity. To resolve this potential issue, users may need to pre-process the genetic data to remove highly correlated genetic markers.

## 4 Results

In this section, we compare the speed, and the maximum memory usage, and accuracy of the original MATLAB (8) and our Python implementations of the ERStruct algorithm. We use the function `tic` in MATLAB and `time.time` in Python to record running time. To record memory usage, we use the Linux shell command `/proc/<pid>/status | grep VmSize` to check memory usage in MATLAB and use the Python module `memory-profiler` for checking in Python. Python (version 3.8.8) is used with NumPy (version 1.20.1), PyTorch (version 1.11.0) and Joblib (version 1.0.1). All results are obtained from a server running x86-64 Linux with 15 Intel(R) Xeon(R) E7-8891 v4 CPU cores and 12 GB Tesla K80 GPU.

We apply our ERStruct Python implementation to the publicly available 1000 Genomes Project (10) data to estimate the number of top informative PCs. The 1000 Genomes Project is a whole genome sequencing dataset with 2504 individuals from 26 subpopulations. Following the same procedure in (8), the raw sequencing data file is first filtered out markers with MAF (Minor Allele Frequency) less than 0.05, 0.01, 0.005, and 0.001 using the PLINK software. The remaining number of markers are *p*_0.05_ = 7, 921, 8816, *p*_0.01_ = 13, 650, 478, *p*_0.005_ = 17, 307, 567 and *p*_0.001_ = 28, 793, 505, respectively. Each preprocessed data is stored as NPY files and tested using our ERStruct Python package, and the original MATLAB toolbox by (8). The testing parameters are fixed as: number of replications `rep` = 5000, significance level *α* = 10^−4^.

The results are shown in Table 1. The GPU-based Python implementation runs much faster than the CPU-based Python (2.02 times faster when MAF is greater than 0.001) and MATLAB (3.78 times faster when MAF is greater than 0.001) implementations. In terms of the maximum memory usage, the GPU-based Python implementation used only 0.27 of the CPU-based Python implementation and 0.31 of the MATLAB implementation (when MAF is greater than 0.001) due to the data splitting procedure. Noting that our function `Eigens` automatically splits large-scale datasets so that our algorithm can be run on a GPU with limited VRAM, so using GPU acceleration can be slower than using CPU only when the current available VRAM is too small. In our testing environment, this only happens when the available VRAM is less than 0.17 GB. It is only possible when another process in the working environment has taken up a large part of the available VRAM. In this case, we suggest releasing the VRAM first. In the package, we set the default VRAM value to 0.2 GB to ensure better performance.

**Table 1:** Running time (in minutes) and maximum memory usage (in GB) comparisons of the ERStruct algorithm, using MATLAB, Python CPU and Python GPU implementations on the 1000 Genomes Project data with different MAF filtering thresholds.

| MAF filtering thresholds |  | 0.05 | 0.01 | 0.005 | 0.001 |
| --- | --- | --- | --- | --- | --- |
| Time | MATLAB | 30.08 | 42.30 | 50.05 | 73.41 |
|  | Python CPU | 34.89 | 66.54 | 84.97 | 137.54 |
|  | Python GPU | 17.60 | 21.91 | 28.96 | 36.42 |
| Memory | MATLAB | 51.55 | 62.22 | 69.54 | 92.50 |
|  | Python CPU | 28.00 | 48.80 | 62.10 | 104.71 |
|  | Python GPU | 10.08 | 15.07 | 18.33 | 28.67 |

To evaluate the accuracy of our ERStruct algorithm implemented in both of the Python package and MATLAB toolbox, we conducted 30 identical experiments on CPU and GPU, respectively. We used the 1000 Genomes dataset with MAF greater than 0.001, and set the number of replications to `rep` = 5000 and the significance level to *α* = 10^−3^. In all 30 experiments, the output consistently estimated 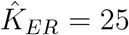. These results indicate that the ERStruct algorithm implemented in both versions are identical and produce the same output.

## 5 Conclusion

In this paper, we developed a Python package that employs the ERStruct algorithm to determine the optimal number of top informative PCs in WGS data. By leveraging parallel computing on CPUs and GPU acceleration, our package demonstrates exceptional efficiency in performing matrix operations on large-scale sequencing datasets. To enhance the usability of our Python package across various environments, it features adaptive data splitting capabilities for GPU computation with limited VRAM. We conducted experiments that compared the computation speed of our ERStruct Python package to the ERStruct MATLAB toolbox in (8). Our results demonstrate a significant improvement in computation speed with the ERStruct Python package.

## Availability

Our Python implementation of the ERStruct Algorithm is freely available at

https://github.com/ecielyang/ERStruct.

The datasets were derived from sources in the public domain:

https://www.internationalgenome.org/data.

## Acknowledgements

The authors thank The University of Hong Kong for providing the computing resources.

## Funding

Dr. Gao Wang is supported by grant R01 AG076901.

## Notes

https://www.internationalgenome.org/data

### Competing Interest Statement

The authors have declared no competing interest.

### Summary of Updates

We enriched the content and included additional references to support our claims. We provided more details about the figure to make it easier to understand.

